# ChemEM: flexible docking of small molecules in Cryo-EM structures using difference maps

**DOI:** 10.1101/2023.03.13.532279

**Authors:** Aaron Sweeney, Thomas Mulvaney, Maya Topf

## Abstract

The rapid advancement of the “resolution revolution” has propelled cryo-electron microscopy (cryo-EM) to the forefront of structure-based drug discovery. However, the majority of cryo-EM structures are solved at medium resolution (3-4Å), an unexplored territory for small-molecule docking, due to difficulty in positioning ligands and the surrounding side-chains. Therefore, the development of software capable of reliably and automatically docking ligands into cryo-EM maps at such resolutions is of utmost importance. ChemEM is a novel method that employs cryo-EM data, difference mapping, and a physico-chemical scoring function to flexibly dock one or multiple ligands in a protein binding site. To validate its effectiveness, ChemEM was assessed using a highly curated benchmark containing 33 experimental cryo-EM structures, spanning a resolution range of 2.2-5.6 Å. In all but one case, the method placed the ligands in the density in an accurate conformation, often better than the PDB deposited solutions. Even without the use of cryo-EM density, the ChemEM scoring function outperformed the well-established docking software AutoDock Vina. Furthermore, the study demonstrates that useful information is present in the map even at low resolutions. ChemEM unlocks the potential of medium-resolution cryo-EM structures for drug discovery.

## Introduction

Advancements in cryo-EM have increased the relevance of the technique in structure-based drug discovery. This is evidenced by an increase in the number of cryo-EM maps deposited in the Electron Microscopy Databank (EMDB)(1) at a resolution of 3Å or better, up from 15 in 2015 to over 2000 in 2022. Advances in the resolution limits of cryo-EM have facilitated the identification of novel drug targets, including membrane proteins (2), which are difficult to solve with other techniques such as X-ray crystallography.

Although atomic resolution is achievable (3), the majority of maps are still deposited with a global resolution between 3 to 4Å (1). Furthermore, most cryo-EM maps contain heterogeneous resolutions across the map. Even in high-resolution maps, small perturbations in the position of ligands, multiple conformations or partial occupancy within the binding site may worsen the local resolution of the binding site relative to the global resolution. Therefore, it is necessary to develop methodologies for the accurate modelling of small molecules at both high- and low-resolution.

Only recently have methodologies that specifically focus on small molecule fitting in cryo-EM emerged (4, 5). The success of such a process is dependent on the initial fit of the ligand, as having a ligand positioned close to the correct position increases the chance of generating a high-quality fit to the map. These methodologies have leveraged molecular docking for generating initial fits. Molecular docking software use scoring functions to describe the interactions between a protein and small molecule and identify physico-chemically reasonable ligand conformations within a given protein binding site. One of the most common softwares, AutoDock Vina (6), was shown to have amongst the best scoring functions for estimating affinity and identifying correct conformations within a set of decoys in the Critical Assessment of Scoring Functions 2016 report (7) (CASF-2016). In the commercial software GemSpot (4) such docking score is integrated with a goodness-of-fit term – the Cross Correlation Coefficient (CCC). This allows only ligand conformations with a good fit to the map and favourable docking score to be selected for further refinement. Whilst the CCC is the *‘gold standard’* goodness-of-fit score for fitting atomic structures in cryo-EM, there is evidence that the Mutual Information (MI) score provides a comparable measure of fit at high-resolution and is more robust to a decrease in resolution when fitting protein models to maps(8). However, data are lacking as to whether this can be applied to small molecule fitting.

Thus far, the goodness-of-fit has been assessed mostly between a density map simulated from the model and the experimental cryo-EM density map. However, methodologies where density-difference maps are used instead of the full map have been employed. For example to fit the BTB-1 ligand bound to a human kinesin-8 motor domain at 4.8Å resolution (9) and the GSK-1 ligand bound to Eg-5 at 3.8Å resolution (10), the molecular docking software GOLD (11), HADDOCK (12), and AutoDock Vina (6) were used along with difference maps for ligand fitting.

Recently, a technique for difference mapping based on local scaling of the cryo-EM map amplitudes was shown to be capable of identifying ligand density by subtracting an apo-protein density map from a map of the protein ligand complex (13). However, there is no data that compares these procedures to using the full map. Additionally, an integrative automated workflow optimised for docking small molecules utilising density difference maps has not yet been proposed.

In the following paper, we describe ChemEM – a novel methodology for docking small molecules into medium-to-high-resolution cryo-EM structures. We first describe a novel molecular docking scoring function (ChemDock) shown to have a performance comparable to or better than the AutoDock Vina scoring function. We then show the usefulness of the MI score in fitting small molecules into cryo-EM difference maps compared to the commonly-used CCC, especially for medium resolutions. Finally, we introduce ChemEM - a fully automated flexible docking software, which combines both ChemDock and the MI score to better identify and refine ligand conformation in binding sites of cryo-EM structures. ChemEM was validated using 32 experimental structures of proteins bound to small molecules at resolutions between 2.2Å and 5.6Å.

## Results

### The ChemEM workflow

ChemEM docks small molecules to cryo-EM maps using a two stage approach (Figure 1). In Stage 1 an approximate fit of a ligand into the protein structure is generated using a difference density map between the protein-ligand complex map and a simulated protein map.

**Figure 1.**
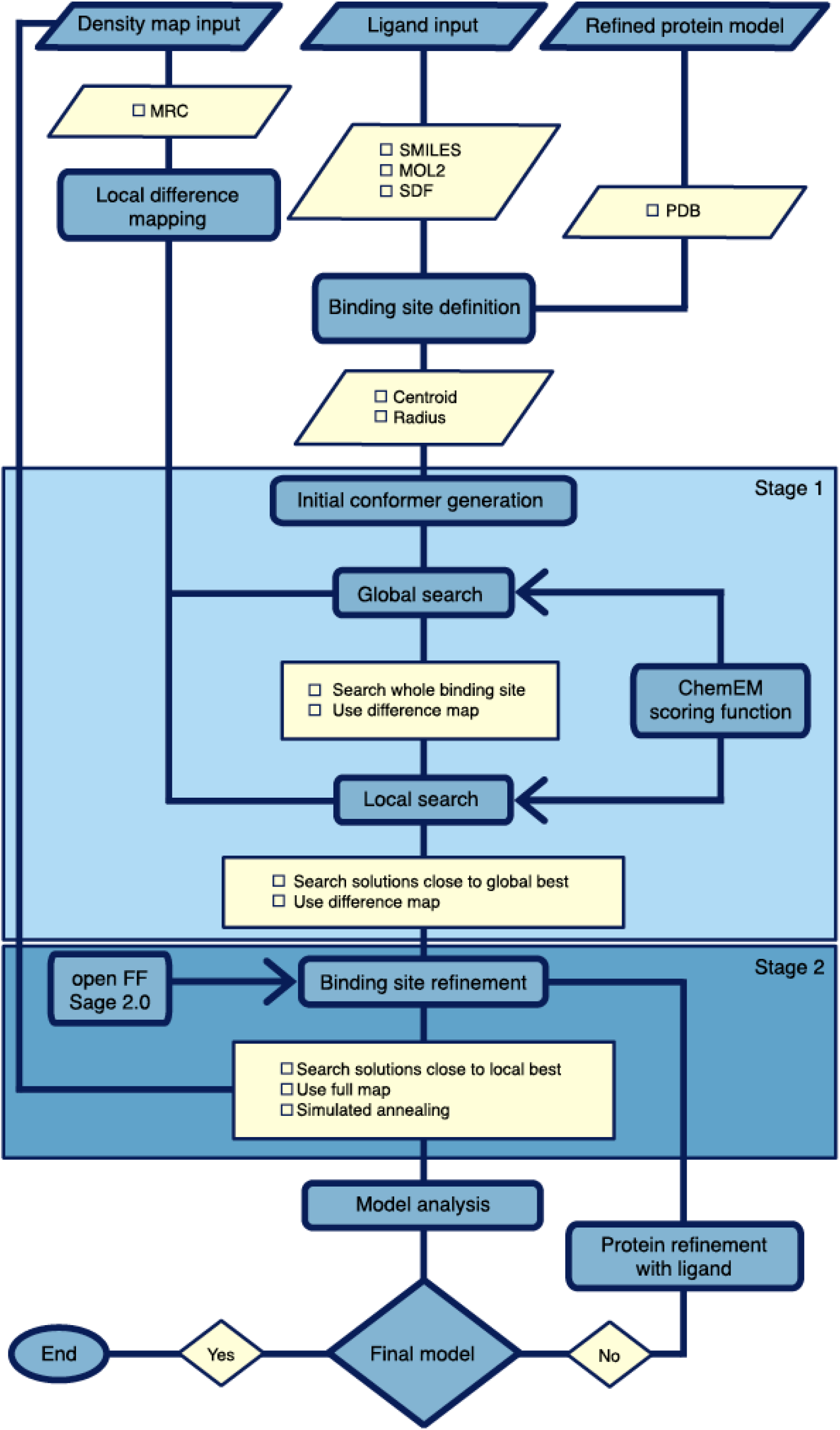
Schematic overview of the ChemEM methodology for docking small molecules, also in combination with cryo-EM maps. For running docking with ChemDock a ligand and protein input are needed. For running ChemEM an additional density map input is required. Additionally, the binding site centroid and radius need to be set by the user to identify protein residues that line the binding site. For ChemEM a local density difference map is created using the user-supplied binding site definitions. For both ChemDock and ChemEM stage 1 generates initial fits of small molecules within the binding site using either the ChemDock or ChemEM scoring functions. The global search is an Ant colony optimisation (ACO) algorithm whilst the local search is a Nelder-Mead minimisation algorithm. Stage 2 refines the initial fits in the context of the binding site. Here, all binding site residue side chains are treated as fully flexible. The openFF Sage v2.0 forcefield is used for small molecule parameterisation, with protein parameters taken from the AMBER FF14SB forcefield. For ChemDock workflow only energy minimisation is applied. For the ChemEM workflow, the full density map term is added to the scoring function and a simulated annealing search algorithm is used to identify final solutions.

An initial fit is achieved by molecular docking using an empirical scoring function integrated with the MI score. This ensures that the generated conformation is both physico-chemically acceptable and well-fitted to the difference map. In this stage, the difference map acts as a constraint for the docking algorithm, reducing the conformational space to be searched.

In Stage 2 (Figure 1), candidate conformations are refined into the full cryo-EM density map using a flexible fitting approach with the AMBER forcefield-14SB (14) protein parameters and the SAGE OpenForceField 2.0.0 (15) parameters for small molecules. This stage fine-tunes the initial fits generated in Stage 1, whilst simultaneously refining the fit of binding-site atoms.

### Evaluation of the ChemDock scoring function

The density independent part of the molecular docking scoring function (termed the ChemDock score) for the ChemEM scoring function included terms for hydrogen bonding, hydrophobic interactions, π-π stacks, steric and repulsive terms (see methods). The score was trained to estimate the pKd of a given protein/ligand complex using 3,281 complexes taken from the PDBBind dataset (Table S1, Table S2).

The correlation between the ChemDock or AutoDock Vina (6) scoring functions and binding affinity was assessed using a benchmark of 153 protein-ligand complexes, each associated with an experimentally determined binding affinity. The predicted binding affinities based on ChemEM had a Pearson correlation of 0.71 to the experimental binding affinities compared to 0.62 for AutoDock Vina.

We next looked at the ability of ChemDock to identify a *correct* conformation given a decoy set and rank decoys given their root-mean-square deviation (RMSD) from the *reference* structure (the PDB deposited structure). For each of the 153 protein-ligand complexes, up to 100 ligand decoys from CASF-2016 dataset (with increasing RMSD values from the reference structure), were ranked by both ChemDock or AutoDock vina scoring functions. ChemDock was able to rank a correct conformation (defined as ≤2.0Å from the reference conformation) within the top 3 conformations in 94.8% of cases, and of these a correct pose was ranked within the top 2 in 86.9%, and top 79.1% of the time. The performance of ChemDock was comparable to that of Vina, where Vina ranked a correct conformation in the top 3rd, 2nd or 1st ranks in 94.12, 90.2, and 79.74% cases, respectively (Figure 2A).

**Figure 2.**
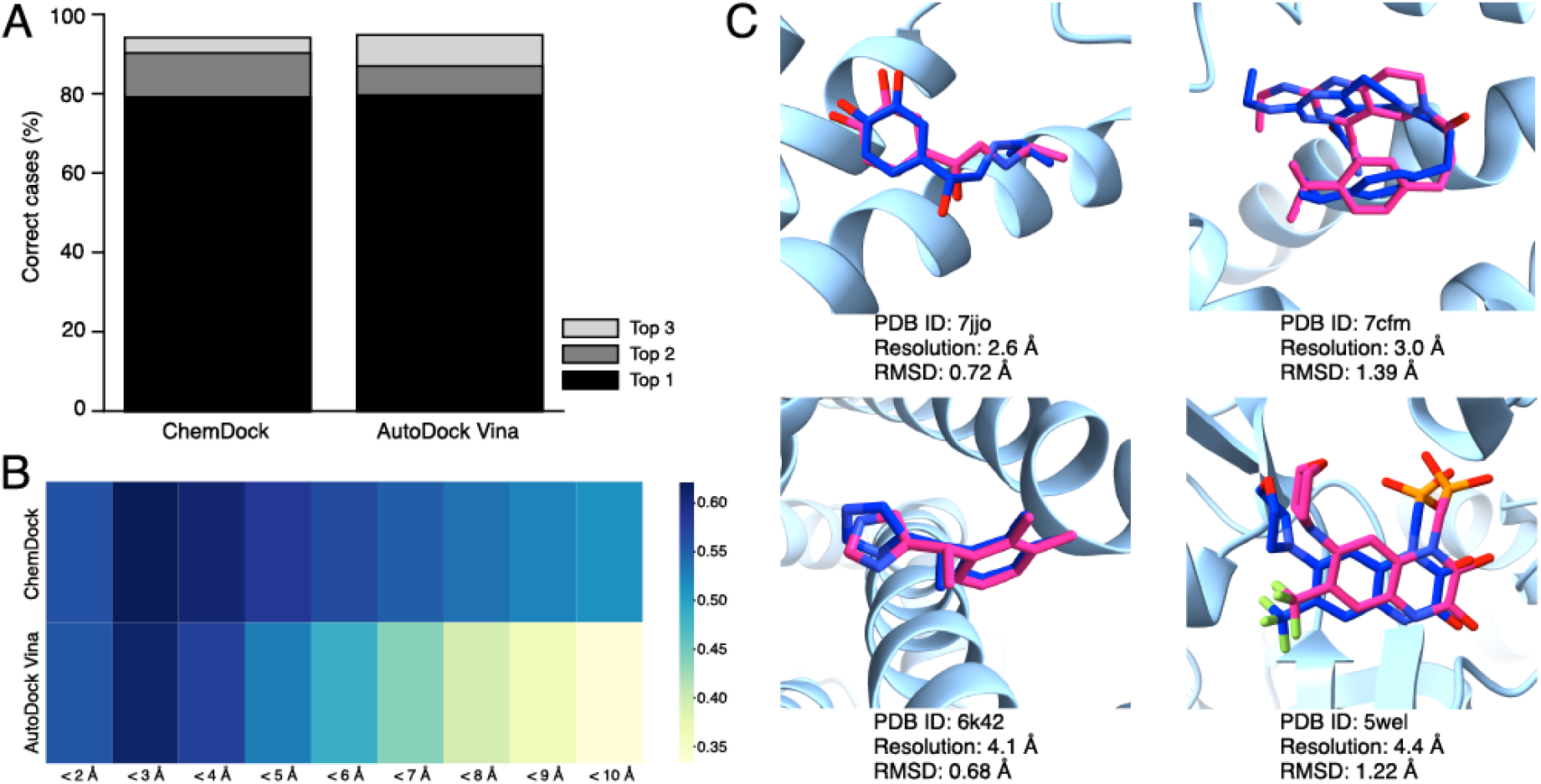
A comparison of the ChemDock score with the Autodock Vina docking score. **A**. The percentage of cases where a conformation was ranked in the top 1 (black), 2 (dark grey), 3 (light grey) positions by both scoring functions. **B.** The binding funnel analysis for both the ChemDock and AutoDock Vina scores. The average Spearman correlation between the RMSD and either scoring function when only solutions < 2, 3, 4, 5, 6, 7, 8, 9, or 10Å RMSD were included in the RMSD-decoy benchmark. The corresponding correlation colour key is shown to the right. **C**. ChemDock solutions (Pink) for cases from the high-resolution benchmark (PDB: 7JJO, 7CFM) and the low-resolution benchmark (PDB: 6K42, 5WEL). The PDB deposited solutions are also shown (dark blue) along with protein binding site elements (light blue). The resolutions and RMSD between solutions are also indicated.

In order to further probe the ChemDock scoring function, the binding funnel analysis introduced in Critical Assessment of Scoring Functions-2016 (7) (CASF-2016) report (see Methods) was utilised. The analysis showed a good correlation between the ranking of poses based on ChemDock and Vina scoring functions within a given RMSD threshold from the reference pose, especially for poses within < 3.0Å threshold (Figure 2B). However, the ChemDock correlation was more robust than that of Vina further away from the reference pose (~6-10Å) (Figure 2).

### Docking small molecules with ChemDock scoring function

To run docking experiments the ChemDock scoring function was integrated with an Ant-Colony Optimisation (ACO) algorithm (16). A ‘high-resolution’ benchmark was constructed, consisting of 20 PDB protein-ligand complexes derived from cryo-EM experiments at 2.2-3.0 Å resolution (Table S3). All deposited ligands were removed from the structures and re-docked starting from a random conformation in the same binding site using the ChemDock software. A correct conformation (< 2Å from the reference conformation) was generated 85% of the time, a *good* conformation (< 1.5Å) in 70% of cases and *excellent* conformation (< 1.0Å) in 15% of cases (Figure 3A). When the same ligands were docked with Autodock Vina, a correct conformation was found in 70% of cases, good in 60% and excellent in 25%. The quality of the final solutions was analysed using the conformational torsional strain energy (17) (a measurement of the quality of conformations). Overall the solutions generated with ChemDock were seen to be of better quality than those generated with Autodock Vina, with a mean minimum RMSD of the top solution (over the entire benchmark) and average torsional energy units (TEU) of 1.44Å and 5.90, compared to 1.93Å and 6.53 (Table S4).

**Figure 3.**
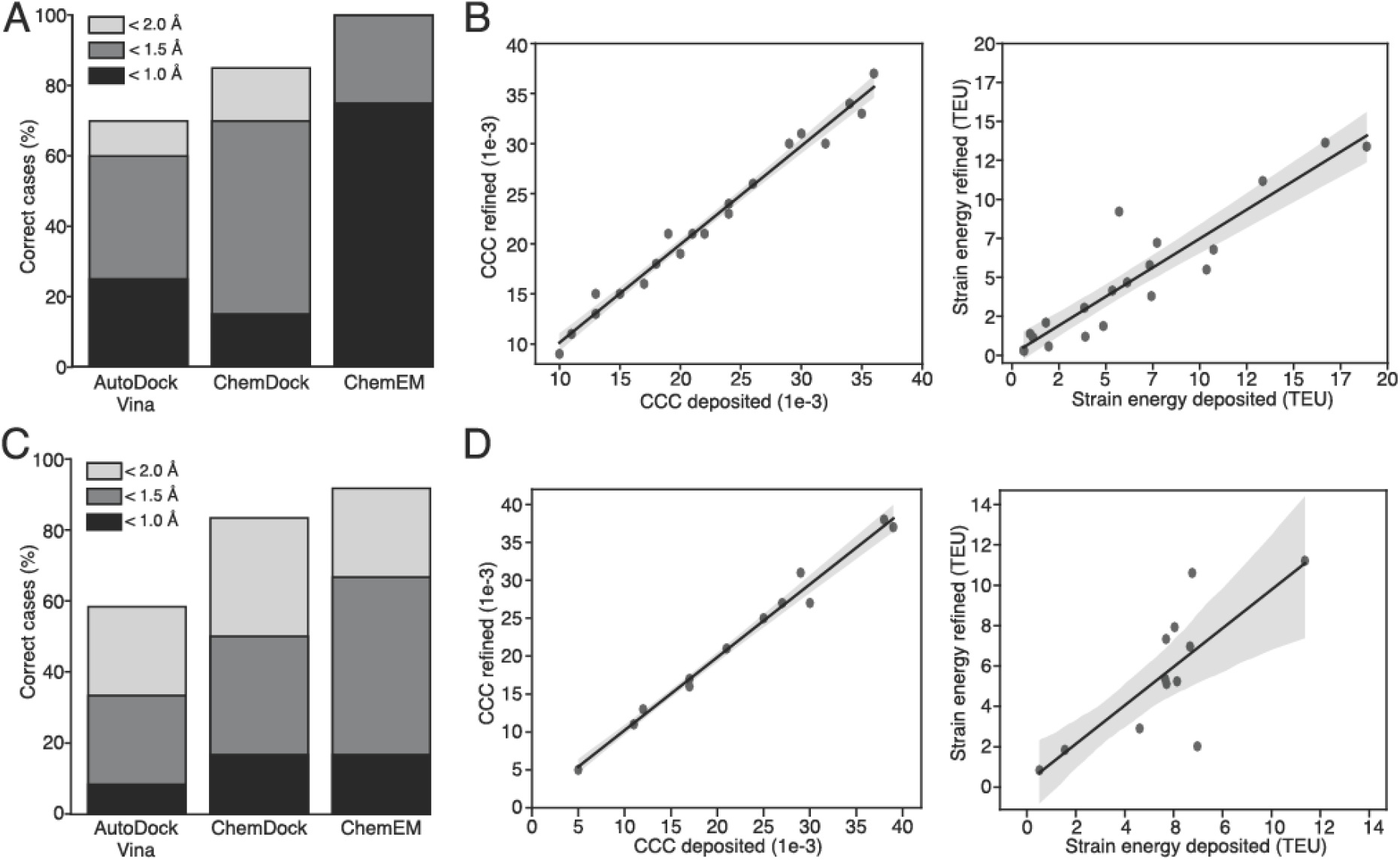
A statistical overview of docking the high- and low-resolution benchmarks with AutoDock Vina, ChemDock and ChemEM. **A**. The percentage of cases from the high-resolution benchmark where docking/fitting produced a correct (< 2Å RMSD from the deposited solution), good (< 1.5Å), or excellent (< 1.0Å) solution. **B.** The correlation between the CCC (left) and the strain energy (right) of the deposited solutions with that of the final ChemEM refined solutions. **C**. The percentage of cases from the low-resolution benchmark where docking/fitting produced a correct, good, or excellent solution. **D.** The low-resolution benchmark correlations between deposited and ChemEM refined solutions for the CCC (right) and strain energy (left).

To test the accuracy of the ChemDock software on a wider range of structures, docking experiments were repeated using 12 structures from lower-resolution cryo-EM maps (3.0 - 4.5Å, Table S3). ChemDock generated a correct conformation in 83% of cases, while good and excellent solutions were found 50% and 16% of the time, respectively (Figure 3C). Autodock Vina generated a correct conformation in 58% of cases, with 33% good and 8% excellent. The mean minimum RMSD from the deposited solution was 1.72Å for ChemEM and 2.55Å for Autodock Vina. However, the solutions generated by autodock vina showed a better mean strain energy of 7.68 TEU compared to 8.32 TEU for ChemEM (Table S5).

Finally, we assessed the ability of ChemDock to rank generated ligand solutions. We found that the rank of the ChemDock solution with the lowest RMSD to the reference solution occurred on average within the top 15% of generated solutions for both high- (Table S6) and low-resolution (Table S7) benchmarks.

### Integrating ChemDock with cryo-EM density (ChemEM)

We investigated the performance of two goodness-of-fit metrics, CCC and MI, for fitting small molecules using a simulated benchmark. Density difference maps were simulated from the X-ray structure corresponding to 285 protein/ligand complexes from the CASF-2016 database (7) at a 2.5-8.5Å resolution range (every 0.5Å). For each case, the CASF-2016 decoy set was ranked by either MI or CCC.

At each resolution tested, for both MI and CCC there was a strong negative correlation between the RMSD and goodness-of-fit metric (Table S8); however, MI had a higher correlation. This difference was statistically significant at the 4.5-8.5Å range. The Pearson correlation coefficient for the MI score remained relatively consistent for each resolution while decreasing for the CCC. This indicated that the MI score was more robust to changes in the resolutions of the difference density maps, inline with previous reports using the MI score to fit protein structures (8).

Next, the ChemEM workflow was tested for fitting small molecules into the above-mentioned experimental cryo-EM maps at both high and low-resolutions. The first part of the workflow fits small molecules to the experimental difference density maps using the ACO algorithm (16, 18). To identify an optimal weight to integrate the MI score with the molecular docking score this step was performed without minimisation. Different MI weights were tested of 10, 50 and 100 that scaled the MI score to 0.1x, 0.5x, and 1x the magnitude of the molecular docking score, respectively.

Using the high-resolution benchmark showed the docking score alone identified a correct ligand conformation in 75% of cases, whilst the MI score alone identified a correct conformation in only 55% of cases (Figure 4A). The success rate of this initial fit rose to 85% when MI was integrated with the docking score at a weight of 50 (Figure 4A). This was accompanied by an increase in solution accuracy compared to using docking or MI scores alone, with 60% of cases generating a solution ≤1.5Å and 25% within 1Å of the deposited solutions (Figure 4A).

**Figure 4.**
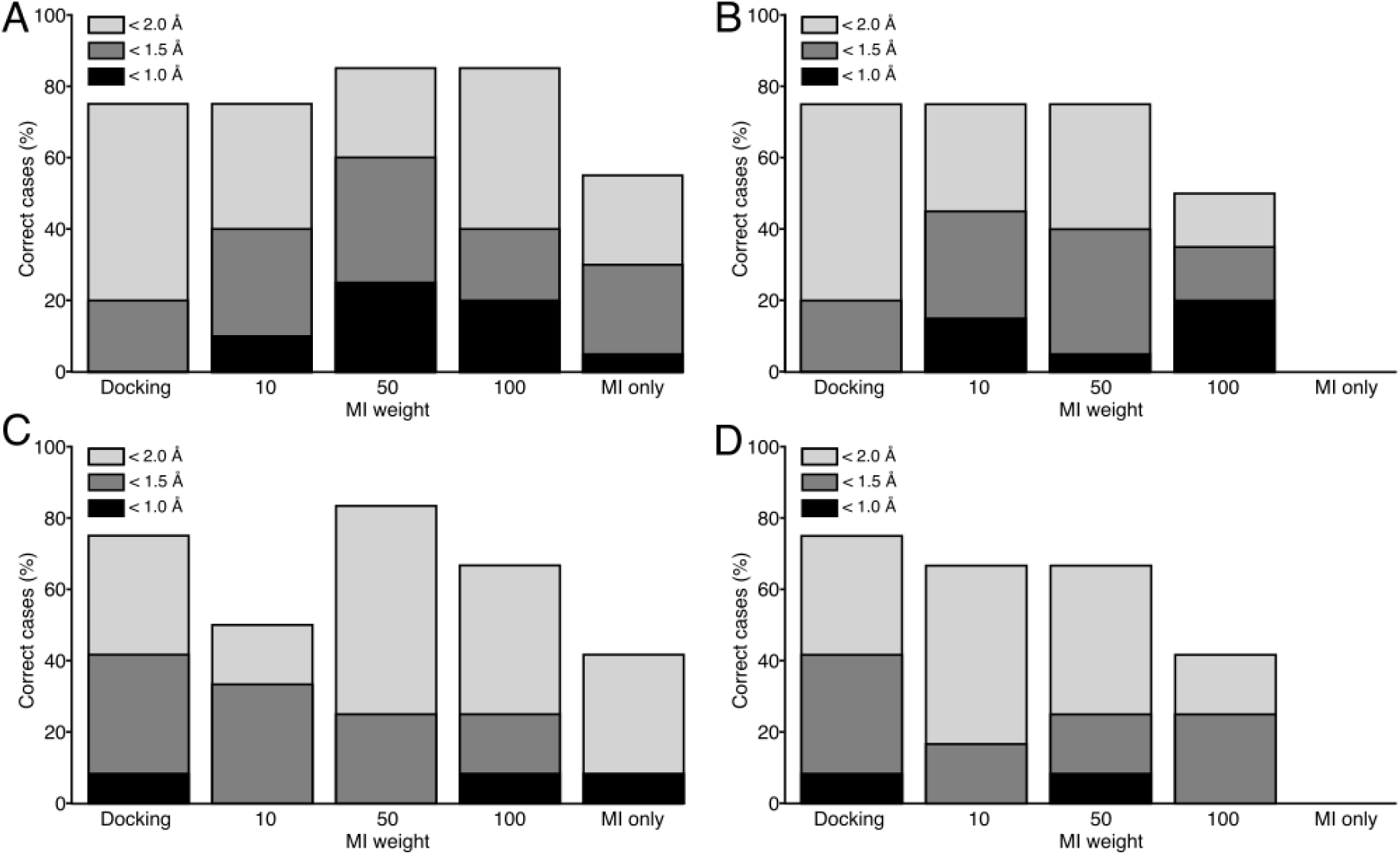
The results of fitting with the ACO and the ChemDock score (docking), MI alone, or the ChemEM score with MI score weights of 10, 50 and 100. **A.** The number of correct cases where a correct solution was generated when the analysis was run the high-resolution benchmark using density difference maps. **B.** The correct cases when the high-resolution and full maps were used. **C.** The number of cases where a correct solution was generated when the analysis was run the low-resolution benchmark using density difference maps. **D**. The correct cases using the full maps and low-resolution benchmark.

To ascertain whether the MI weight of 50 was applicable at resolutions >3.0Å, the experiment was repeated using the lower resolution experimental benchmark. Once again a combination of the MI and docking score identified the most correct solutions (83%) compared to each score alone (75% and 42% correct cases, respectively) (Figure 4C). However, only 25% were identified at an RMSD of ≤1.5Å from the deposited ligand.

Assessing the rank of the ChemEM solution with the lowest RMSD to the reference solution showed that the solution with the lowest RMSD appeared in the top 5% of solutions on average for the high-resolution benchmark (Table S9) and the top 10% for the low-resolution benchmark (Table S10).

Finally, the ChemEM scoring function performed better using difference maps compared to full maps. For the high-resolution benchmark, no MI weight was found to generate more correct solutions using full maps compared to docking alone (Figure 4B). However, integrating MI did improve the solution quality, resulting in 45% good and 15% excellent solutions with MI weight of 10 (Figure 4B). With lower resolutions, using full maps resulted in a lower number of correct solutions when integrating MI (compared to docking alone), with the best MI weights (10 and 50) resulting in only 66% of cases having a correct solution. Furthermore, the solutions were of a lower overall quality compared to docking alone (Figure 4D).

### Flexible fitting to full maps

For each case in the high- and low-resolution benchmark the best initial fits were generated using density difference maps and the ChemEM scoring function with an MI weight of 50. Following this molecular docking protocol, the solutions were further refined into the full map with a flexible fitting protocol (Methods, Figure 1).

In all 20 cases of the high-resolution benchmark a good solution was generated, with an excellent solution found 75% of the time (Figure 3A). The CCC of the refined solutions (in the original full map) correlated well with that of the reference solutions, with a mean CCC of 0.023±0.008 STD for the reference conformations and 0.022±0.008 STD for the refined solutions (Figure 3B). However, the quality of the conformations generated by ChemEM was notably better than that of the deposited solutions with mean strain energies of 4.86 and 6.47 TEU, respectively (Figure 3B).

The workflow was repeated using the 12 low-resolution protein-ligand complexes. Following four rounds of flexible fitting, 92% of solutions were correct (Figure 3C). This was accompanied by an increase in the quality of solutions with 67% good solutions and 17% excellent. The mean CCC was 0.022±0.01 and 0.023±0.01 for the refined and reference conformations, respectively (Figure 3D). The strain energies over the whole dataset were inline with the published structures with a mean of 5.61 TEU compared to 5.64 and 5.99 TEU in the reference structures and high-resolution controls, respectively (Figure 3D, Table S5).

### Analysis of specific examples

#### High-resolution benchmark (≤ 3.0)

As discussed above, the ChemEM software was able to find meaningful solutions for most of the 20 high-resolution structures that agreed well with the deposited ligands. In a few cases, using ChemEM led to only modest improvements in the RMSD from the reference pose compared to docking alone. However, in most cases it led to much-improved fits, with a mean CCC of 0.022±0.008 for the best solutions, compared to 0.019±0.007 using docking alone (Table S4). Additionally, the mean ligand strain energies decreased from 8.12 to 4.86 TEU (Table S4). For example, in the case of Isoprenaline bound to an avian ***β***-Adrenergic receptor protein (19) (PDB: 7JJO, EMD: 22357) the docking solution had an RMSD of 0.72Å compared to 0.64Å when the MI score and flexible fitting protocol were included (Figure 5A).

**Figure 5.**
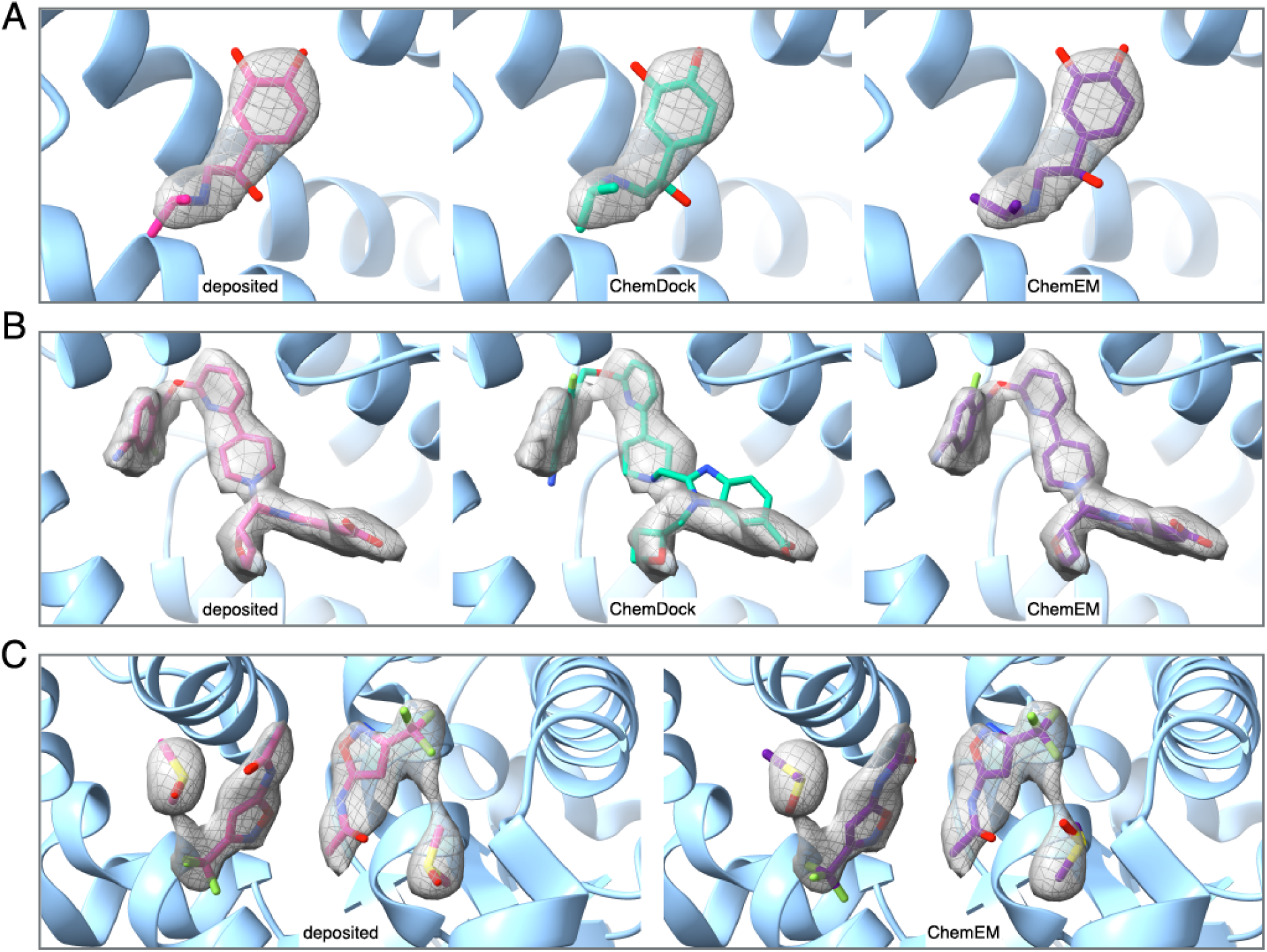
Specific examples of ChemDock (Docking) and ChemEM (refined) solutions for the high-resolution benchmark compared to the deposited solutions when: **A.** Isoprenaline was fit to the ***β***-Adrenergic receptor protein (PDB: 7JJO, EMD: 22357). **B.** The small molecule PF06882961 bound to the human glucagon-like peptide-1 receptor (PDB: 6X1A, EMD: 21994) **C.** Two fragment molecules and two molecules of dimethyl sulfoxide (DMSO) were fitted to the human pyruvate kinase protein (PDB: 6TTI, EMD: 10577).

However, in the case of PF06882961 bound to the human glucagon-like peptide-1 receptor (20) (PDB: 6X1A, EMD: 21994), docking with ChemDock failed to find a correct ligand conformation, with RMSD of2.09Å from the reference pose for the best solution (Figure 5B), a CCC of 0.017 and a strain energy of 13.41 TEU, compared with a CCC and strain energy of 0.032 and 13.38 TEU for the reference structure. When the cryo-EM map was used for fitting, ChemEM was able to generate a correct solution with an RMSD, CCC and strain energy of 0.89Å, 0.03 and 11.18 TEU (Figure 5B).

#### *low-resolution benchmark* (> 3.0 - 4.5Å)

For the 12 lower-resolution complexes, the docking software with ChemDock score was also able to find meaningful solutions within the top scored solutions that agreed well with the reference ligands. Similar to the high-resolution cases, using the full ChemEM protocol (MI and flexible fitting) generally led to improvement in the RMSD from the reference pose. An example of the benefit of including the MI score was in the case of CGP 54626 bound to the GABA_B(21)_ receptor at 3.52Å (PDB: 7CUM, EMD: 30472), where docking alone was not able to yield a correct solution (Figure 6A). The full ChemEM protocol improved the fit significantly, resulting in a conformation close to the reference conformation and markedly better fit to the density than the cryoEM deposited structure (Figure 6A).

**Figure 6.**
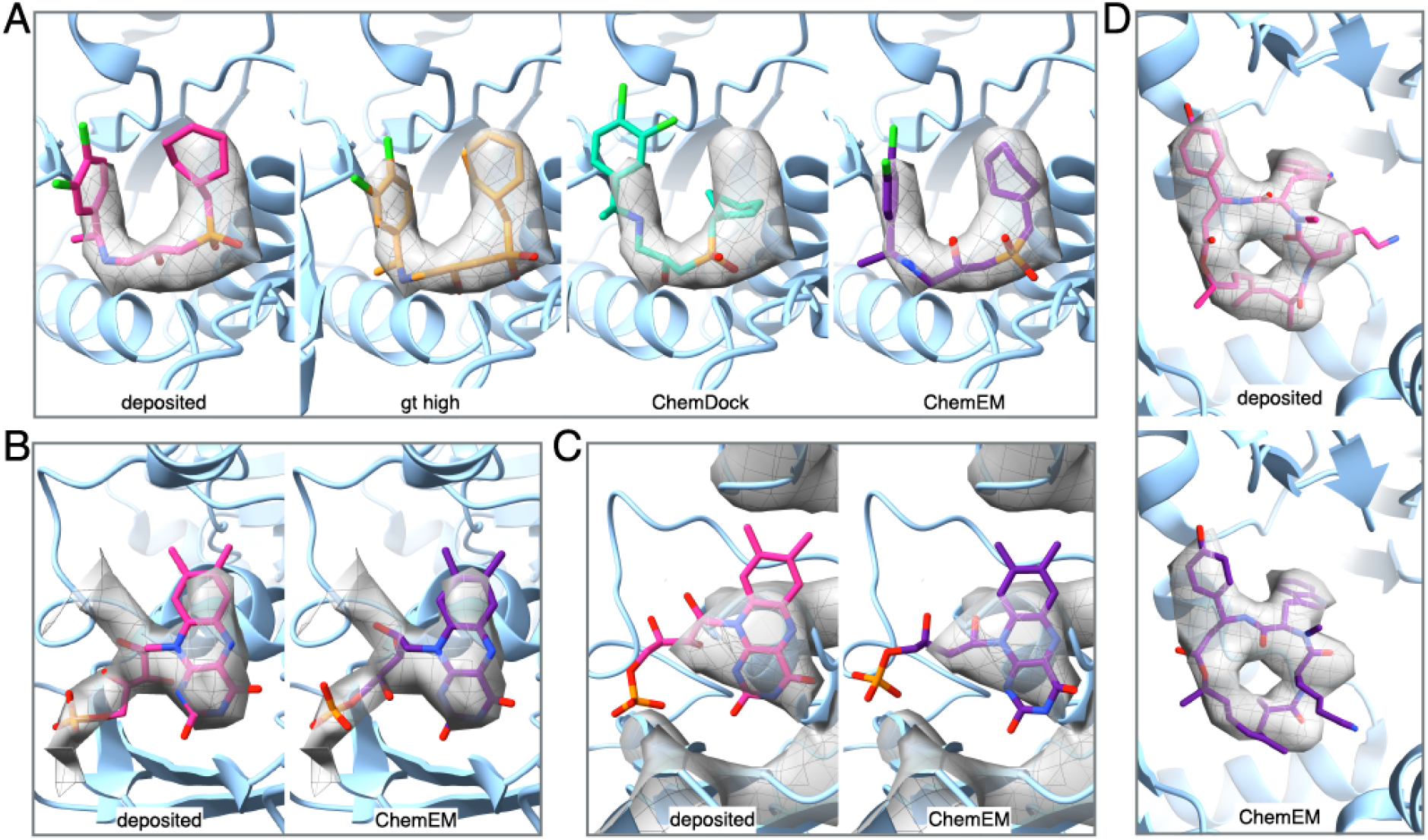
Specific examples of ChemDock (Docking) and ChemEM (refined) solutions for the low-resolution benchmark compared to the deposited and ground truth solutions when: **A.** the molecule CGP 54626 was fit to Human GABAB (PDB: 7CUM, EMD: 30472). **B.** A molecule of Flavin mononucleotide was fit to the bacterial respiratory complex I (PDB: 6ZIY, EMD: 11231). **C.** Jasplakinolide was fit to actin (PDB: 6T24, EMD: 10366).

The ChemEM workflow produced accurate fits even at the lower end of the resolution range tested. One example was the case of Flavin mononucleotide bound to the bacterial respiratory complex I (22) (PDB: 6ZIY, EMD: 11231) at 4.25Å resolution. Following refinement, the CCC was 0.005, equal to that of the deposited solution, along with a strain energy better than the high-resolution control structure and comparable to the reference ligand (Figure 6B). The workflow was only systematically tested on lower-resolution structures at 3.00-4.5Å resolution range, as at worse resolutions it was difficult to identify structures with ligands that had high-resolution control structures deposited in the PDB. However, one interesting case to test was a homolog of the above structure – the Flavin mononucleotide bound to the mammalian respiratory complex I (23) at 5.6Å (PDB: 5LDX, EMD: 4041). At this resolution it was still possible to fit the ligand in a correct conformation, although it was necessary to reduce the MI weight to 10. The final solution had an RMSD of 1.25Å and correlated well with the conformation seen in the reference and high-resolution structures. The CCC and strain energy of the ChemEM solution were 0.023 and 8.06 TEU, notably better than that of the reference solution (CCC: 0.022, strain energy: 14.72, Figure 6C). When the same ligand was fitted using only ChemDock the RMSD of the best solution was 1.71Å, and the CCC and TEU were 0.022 and 12.36, respectively, worse than the solution generated when the cryo-EM map was included.

As described above, the ChemEM methodology resulted in a similar solution to the reference conformations in all cases of the high-resolution benchmark and all-but-one of the low-resolution cases. This refers to the case of Jasplakinolide-bound actin (24) at 3.7Å resolution (PDB: 6T24, EMD: 10366). Jasplakinolide is a large ligand with 47 heavy atoms, and although the aromatic moieties were accurately placed in the density, the large acyl-ring did not converge to the correct solution (Figure 6D).

### Multi ligand fitting

One example within the high-resolution benchmark described the use of the human pyruvate kinase protein (25) (PDB: 6TTI, EMD: 10577) for fragment screening using cryo-EM. The binding site contained multiple ligands, with two “fragment” molecules and two molecules of dimethyl sulfoxide (DMSO).

All four ligands were fitted using the ChemEM workflow (Figure 5C). The final conformations had an average RMSD of 1.38Å from the reference structure, over all four ligands. The individual RMSDs of the fragment molecules were 0.38Å and 1.17Å. In comparison, the two DMSO molecules were less-well fitted, with RMSDs of 1.88Å and 2.09Å.

The CCC across all four ligands was 0.016, slightly lower than the reference (CCC: 0.019). However, for the two fragment molecules, the CCCs were the same (0.011 and 0.009). Furthermore, the torsional strain energies of both the refined fragments and reference fragments were comparable, with TEUs of 1.14 and 1.89, and 1.12 and 1.17, respectively. The fit of the DMSO molecules was less good with CCCs of 0.006 and 0.005, compared to 0.01 and 0.09 in the reference structure, although the two molecules were still placed in the correct region of the map.

## Discussion

There is currently a requirement for a diverse range of software to automatically fit small molecules to cryo-EM structures due to significant developments in this technique over the last decade, making it increasingly relevant to drug discovery. ChemEM is a novel method that meets these needs by combining small-molecule docking with density fitting using difference maps. The method, which can be used with or without the cryoEM data (as a “classical” docking method), was assessed on a benchmark of cryo-EM protein-ligand complexes at 2.2-4.5Å resolution with corresponding X-ray structures. This resolution range is consistent with what is currently required in the field, with 22.5% of maps deposited within the EMDB in 2022 at resolutions better than 3.0Å, and 47.8% between the 3.0 and 4.0Å range. Additionally, ChemEM was able to produce accurate fits for an individual test case at a lower resolution (5.6Å).

The ChemEM ChemDock scoring function was derived to take account of the physico-chemical features of protein-ligand interactions. It was shown to outperform AutoDock vina scoring function when estimating the relative affinity of compounds, and had a comparable performance when identifying a correct ligand conformation from an incorrect one (Figure 2A). Importantly, ChemDock had an improved performance in the binding funnel analysis over AutoDock Vina (Figure 2B). It has been suggested that docking scores that can perform well on this task are good candidates for ‘blind’ docking scoring functions (7).

When ChemDock score was combined with a search algorithm, a correct conformation was generated 85% and 83% of the time for the benchmarks derived from high and low-resolution structures, respectively, a marked improvement compared to AutoDock Vina (Figure 3). Additionally, when both scoring functions failed to reach a correct solution, the solutions generated by ChemDock were generally closer to the reference (Table S4, S5),which was useful when ChemDock was combined with the cryo-EM data.

To achieve this, the cryo-EM density difference map was used, and the MI score was added to the ChemDock score (ChemEM). It has previously been shown that MI-based scores are generally more discriminatory than CCC-based scores for fitting proteins to cryo-EM maps (26, 27). Our results support these findings, showing that when fitting small molecules to density maps at resolutions 2.5-8.5Å, MI has a higher discriminatory power than CCC (Table S8).

When using the cryo-EM density with ChemEM, difference maps helped improve the predictions further than full maps (Figure 4), likely due to the difference map ability to focus the search to the relevant density. Although we have previously used cryo-EM difference maps for ligand docking (9, 10), a systematic comparison with using full density maps has not been undertaken. Our study shows that the advantage of using difference maps at 2.2-3.0Å resolution is reduced when fitting ligands in lower-resolution densities (3.0-4.5Å) and this effect holds true with full maps. This is due to an increased search space, leading to more false positive solutions being identified.

At lower resolution our investigation indicated that more onus should be placed on the physico-chemical properties rather than the goodness-of-fit to derive a good solution. This was exemplified when the ChemEM score was used for fitting without refinement: in the high-resolution benchmark a greater MI weight of 100 resulted in the same number of correct solutions identified with a weight of 50 (Figure 4). Whereas, for the lower resolution benchmark when the contribution of the MI score was increased to 100, the number of correctly identified solutions decreased (Figure 4). Naturally, this was a consequence of less information being present in the map. However, the fact that this effect starts to occur at resolutions close to 3.0Å is somewhat surprising.

Once initial fits are identified with ChemEM (Stage 1, Figure 1), the addition of refinement in the density in combination with the OpenMM sage v2.0.0 forcefield (Stage 2, Figure 1), improved the quality (TEU, RMSD < 1.5/1.0Å) and success rate (RMSD < 2.0Å) of solutions (Figure 3A, 3C). This improvement was most pronounced for the high-resolution benchmark (Table S4). However, even in the low-resolution benchmark, only one case failed to identify a correct solution (Figure 6D), while in another case (PDB: 7CUM, EMD: 30472 (21)) ChemEM identified a correct solution while the conformation in the deposited cryo-EM structure was clearly incorrect (Figure 6A). Additionally, as seen for generating the initial fits with ChemEM, the best results after refinement for the lower resolution benchmark were achieved when the contribution of the map component within the score was lower in comparison to the high-resolution cases (Table S4). One important point to note, is that when fitting into maps at resolutions worse than 3.0Å, having less information present in the map will not only affect the accuracy of the ligand placement in the map but also of the protein side-chain placement. This in turn may affect the density-independent part of the scoring function as misplaced side-chains may lead to improper bonds being scored between protein and ligand. The flexible fitting stage (stage 2, Figure 1) of the ChemEM workflow aims to minimise these errors by allowing the protein side-chains to move during refinement.

Furthermore, adding the density not only increased the number of solutions where a correct case was generated, but also improved their rank. This explains why there was a greater improvement in the ranking of the best solutions in the high-resolution benchmark (Table S6, S9) over the lower-resolution one (Table S7, S10).

In addition to the resolution of the map, the quality of the generated solutions (as assessed by RMSD to the reference solution) is limited by the number of rotatable bonds contained within the ligand (Figure S1). As the number of rotatable bonds increases the RMSD from the reference increases. This effect was seen in both the high- and low-resolution benchmarks and was found to be independent of the resolution. It is most likely a consequence of the increased search space needed to evaluate solutions with a larger number of rotatable bonds. This issue is somewhat taken into consideration within the ChemEM workflow as the number of iterations of the ACO algorithm will increase as a function of the number of rotatable bonds to account for the greater search space needed.

As explained above, our benchmark only cases better than 4.5Å resolution except for a single case at 5.6Å resolution (Figure 6C), where it was possible to use ChemEM to dock a Flavin mononucleotide to the mammalian respiratory complex I, yielding a solution that fits the map better than the PDB-deposited solution (23). While fewer structures at this resolution contain small molecules, ~14% of the total EMDB deposited maps in 2022 were between 4-6Å resolution. At this resolution range a good fit could only be achieved by reducing the MI weight from 50 to 10 in the ChemEM score. However, including the map with low weight was still helpful. This may indicate that at relatively low resolutions there may still be some benefit to including the map in docking small molecules, even though more weight is needed to be placed on forcefield scores.

Finally, the ChemEM software was extended for the automated simultaneous fitting of multiple ligands (Figure 5), which was computationally feasible owing to the use of difference maps. This feature was shown to work relatively well, with 3 of 4 ligands fitted correctly. Future work would extend this feature for the parallel fitting of other binding site components, for example ions.

## Materials and Methods

### Computational Datasets

Data to train the ChemDock scoring function was taken from the PDBBind-v2020 refined dataset (28). This dataset consists of 5316 protein-ligand atomic models derived from X-ray crystallography at resolutions better than 2.5Å. PDB files that failed to load into the software or those that formed part of the CASF-2016 dataset were removed from the training set, leaving 3281 complexes in the final training set.

The dataset for testing the scoring function was taken from the CASF-2016 data set consisting of 285 protein-ligand atomic models from X-ray crystallography experiments, at resolutions better than 2.5Å (7). When comparing the ChemDock scoring function to the Autodock Vina scoring function any files that failed to load into the ChemDock software or the Open Drug Discovery Toolkit (ODDT) software (29) (used for Autodock Vina rescoring) were discarded from the dataset. The final dataset contained 153 protein-ligand complexes.

The investigations of the CCC and MI goodness-of-fit scores used all 285 protein models in the CASF-2016 dataset (7). Additionally, for each case in the CASF-2016 dataset, up to 100 ligand decoys were included. These decoys were ligand conformations obtained from molecular docking softwares that had RMSDs distributed from 0.0 to 10.0Å from the deposited conformation (7). Simulated maps were generated from the deposited protein and ligand models in USCF-Chimera (30) using the *‘molmap’* command. For each complex maps were generated at resolutions from 2.5-8.5Å in 0.5Å increments. Simulated density difference maps were generated between the full map and the deposited protein model (without the ligand) using a local density difference mapping methodology (13) implemented in TEMPy2 (31).

Two experimental datasets were generated, one containing high-resolution structures (≤ 3.0Å, “high-resolution” benchmark) and a second containing lower resolution structures ( 4.5Å ≤ resolution > 3.0Å, “low-resolution” benchmark). For the high-resolution benchmark, 20 protein-ligand complexes from the Protein Data Bank (PDB) (32) that had an associated Cryo-EM map in the Electron Microscopy Databank (EMDB) (1) were selected. Similarly, 12 protein-ligand complexes were selected for the low-resolution benchmark, however, for the low-resolution structures to be included a *high-resolution control* ( ≤ 3.0Å, representing the ground truth structure for experiments conducted with this dataset) structure of the same protein-ligand complex had to be available in the PDB. Experimental density difference maps were generated using the deposited maps and protein models for each case, using the same methodology used for the simulated maps.

The reason we chose not to systematically test cases at worse resolution was that we wanted to have a high-resolution control structure for each low-resolution case (3-4.5Å) and we could not find cases beyond this resolution range. However, one case above this resolution was chosen where a control structure was available (PDB: 5LDX, EMD: 4041) (23).

### ChemDock scoring function

The density-independent docking scoring function was designed to predict the pKd of a given protein-ligand complex. The score contained terms for atom-atom attraction/repulsion, hydrogen bonding, ***π-π*** stacking, hydrophobic interactions and ligand intra-molecular interactions (Eq. 1):

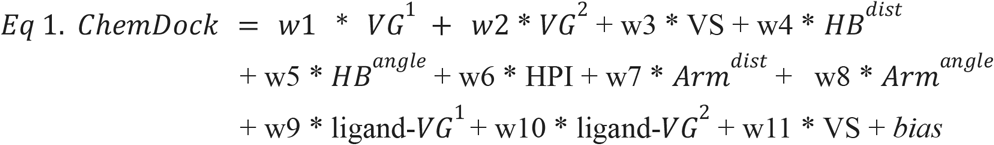

Here we keep the general term for scoring atom-atom steric interactions for Vina(6). This score is the combination of two gaussian functions (Eq. 1, VG^1^, VG^2^) and a repulsion function (Eq. 1, VS). This function was also used to score the intramolecular interactions between non-consecutive atoms of the ligand (Eq. 1, ligand-VG^1^, ligand-VG^2^, ligand-VS).

Hydrogen bonding was accounted for using two geometric parameters, Hbond-Donor-Acceptor distance (Eq 2, HB^dist^) and Hbond-Donor-Hydrogen-Acceptor angle (Eq 3, HB^angle^). The distance score was a function of the van der Waals overlap of the hydrogen-bonded atoms.

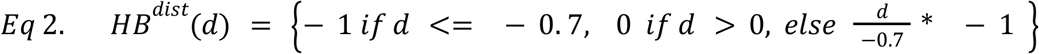

where *d* = *d_ij_* – *r_i_* – *r_j_*;

Where, *d_ij_* is the distance between atomic centres of protein atom *i* and ligand atom *j, r_i_*/*r_j_* is the van der Waals radii of atom *i* or *j*, respectively.

The term to describe the hydrogen bond angle was a linear interpolation between 0 and 1 for hydrogen bond angles between 90° and 180° (Eq 3).

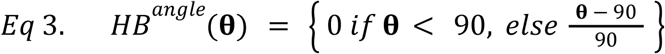

To score hydrophobic interactions the hydrophobic matching algorithm was used, first outlined in SCORE(33). This scored the local environment for a given hydrophobic atom. Hydrophobic atoms were defined as any carbon atoms that were bound exclusively to any carbon atom or hydrogen.

The environment that a hydrophobic atom is placed in was scored as the sum of the logP (a measure of hydrophobicity) of all atoms within 6.0Å of the hydrophobic atom as in X-CSCORE (34) and SCORE (33). Values for logP were taken from XlogP3 (35), however, unlike X-CSCORE and SCORE, the hydrophobic score (Eq 5, HPI) was the sum of the logP values for each hydrophobic atom in the ligand (as opposed to a binary 1 or 0 score if the hydrophobic atom is or is not within a local hydrophobic environment).

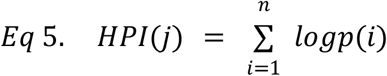

Where *j* is a given hydrophobic atom and *n* is the set of atoms within 6.0Å of atom *j*.

To investigate the preferred geometry of *π-π* stacking interactions the Tough-D1 dataset (36) was used. This data set contained 3079 protein ligand complexes where the ligand was stabilised by at least 1 aromatic ring. For all protein-ligand aromatic *π-π* stacking interactions, data regarding the angle between interacting aromatic rings and the distance between ring centres was extracted. The data was split into two sets, one set containing distance and angle data for *π-π* P-stacks (angle > 45 °) and one for *π-π* T-stacks (angle < 45 °). From this data, the aim was to estimate the probability of a *π-π* stacking interaction being true given the ring-plane angle and ring centre-ring centre distance values.

To do this, probability density functions were derived by fitting the data to 106 common distributions with the python package Fitter. Each fit was scored using the residual sum-of-squares criteria. The distributions that best describe the angle data for both P-stack and T-stack data sets were seen to be beta normal distributions, with sum of the square residuals of 0.0037 and 0.0023, respectively. For the distance data in the P-stack set an exponential normal distribution was seen to fit the data best with the sum of the squared residuals being 0.29, whilst for the T-stack distance data, a skewed normal distribution was seen to fit the data best, where the sum of the squared residuals was seen to be 0.13.

For inclusion in the scoring function the aromatic score was defined as follows:

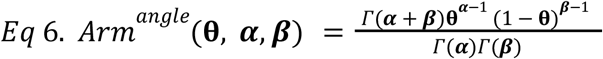

Where ***θ*** is the angle between two planes, ***α*** and ***β*** are constants set based on the angle value, and ***Γ*** is the gamma function: ***Γ***(*n*) = (*n* – 1)!

The score for the aromatic distance was dependent on the plane-plane angle. If the angle was lower than or equal to 45° the interaction was scored using the equation:

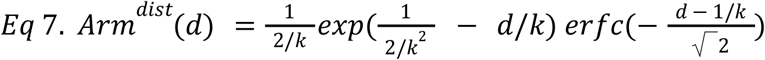

Where *d* was the ring centre-ring centre distance, k is a constant and erfc was the complementary error function. If the angle was greater than 45° the aromatic distance is scored as:

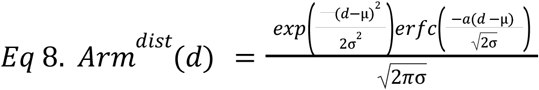

Where, *d* was the ring centre-ring centre distance, σ and μ were constants, and erfc is the complementary error function.

The final ChemDock scoring function was a summation of the weighted terms of all atom-atom pairs within the binding site (Eq 1.). Additionally, a bias term (*bias*) was included to bring the magnitude of the score in line with expected pKd values.

### Fitting the weights in ChemDock scoring function

Each term in the scoring function had to be fitted to a weight. This fitting was done using a linear regression to the experimentally-determined pKd values from the PDBBind-v2020 data set (28).

A 5-fold validation methodology was used for the regression. Briefly, the individual scoring terms were pre-calculated with the ChemEM software for each protein-ligand complex in the final PDBBind-v2020 refined dataset (28). The order of the cases in the dataset was randomised and split into 5 approximately equal sets of each containing 609 complexes. The weights were fit 5 times using 4 of the sets leaving a unique set each time for validating the parameters. The weights for the scores were determined by linear regression to the experimentally determined pKd values (Table S1). Following the linear regression, the weights for each scoring term were averaged from the values in each round (Table S2).

### ChemEM scoring function

When fitting into density maps, the MI score was included in the calculation. When combined with the density independent part of the scoring function, the MI score was weighted by a constant, *w^MI^* (Eq 7.).

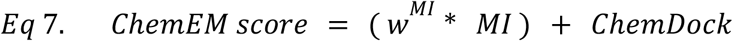

To identify the weight for the MI score in the ChemEM scoring function, docking of both the high and low-resolution benchmarks was conducted in the absence of any further refinement. The weight that gave the best results at both high and low-resolution was seen to be a weight that was approximately 0.5x the magnitude of the density independent part of the scoring function (a weight value of 50, Figure 4). This value was used for all further experiments with the full ChemEM scoring function.

### ChemDock Scoring Evaluation

To evaluate the ChemDock and AutoDock Vina scoring functions the 285 protein ligand complexes in the CASF-2016 benchmark set were used (7). For scoring with ChemDock the supplied protein PDB files were uploaded into the ChemEM software along with the native ligand and each ligand in the associated RMSD benchmark for each case (in MOL2 format). For each protein all benchmark ligand decoys (RMSD benchmark) were scored individually using the ChemDock scoring function with no additional docking applied. To score protein ligand complexes with AutoDock Vina, the Open Drug Discovery Toolkit (ODDT) (29) and an ‘*in house’* python script were used. The ODDT was used as AutoDock Vina does not currently support scoring without docking. Of the 285 cases, 132 failed to load into the ODDT leaving 153 for evaluation.

To evaluate the program’s ability to rank correct ligands, the RMSD benchmark was used. For both ChemDock and AutoDock Vina the solutions in the RMSD benchmark were scored and ranked accordingly. The number of occurrences of either scoring function ranking a correct ligand (defined as < 2.0Å from the deposited conformation) at position 1, 2 or 3 was calculated.

For the binding funnel analysis, the RMSD benchmark was split into subsets containing all the ligands for that case that had an RMSD below a certain cutoff. The cutoffs were slowly raised in steps of 1.0Å from 2.0Å to 10.0Å. At each cutoff, for each case, the subsets were ranked by either the AutoDock Vina or ChemDock score and their respective RMSDs, and the Spearman correlation was calculated.

To evaluate the correlation between the predicted and experimental values a Pearson Correlation Coefficient was calculated between the ChemDock or AutoDock Vina scores for the experimental protein/ligand complex and experimental affinity values supplied with the CASF-2016 benchmark set.

All correlation coefficients were calculated using the SciPy python software and an in-house python script.

### The molecular docking search algorithm

ChemEM and ChemDock used a min-max Ant Colony Optimisation (ACO) Algorithm combined with the Nelder-Mead local minimiser, as in PLANTS (16). The iterations of the search algorithm was proportional to the number of rotatable bonds within a ligand as in (16). When ChemDock was run for molecular docking the output solutions of the ACO were energy minimised within the binding site using OpenMM (37). During this step, all side chains at the surface of the pocket were treated as flexible. Protein parameters were taken from the AMBER FF14SB (14) with small molecule parameters from the OpenFF Sage 2.0.0 (15). Minimization was conducted with the *‘minimiseEnergy’* function in OpenMM with the tolerance set to 10 kJ/mol and allowed to run until convergence. For fitting small molecules in cryo-EM density maps, the algorithm was sped up by segmenting the supplied density map around the centre of the binding site, into a cubic map containing all voxels within a specified distance (30Å by default). This segmented map was used to generate the difference maps for molecular docking (see above).

Docking solutions were further refined using simulated annealing into the segmented full map. Once again both the AMBER FF14SB (14) and Sage OpenFF 2.0.0 (15) were used. Additionally, the full map was added to the potential (Eq 8.) biassing atom movements towards areas of high density. The weight of the map potential was controlled by a constant *k*. Here, experiments were conducted using 3 values of *k*, 25, 50, and 75 (Table S11).

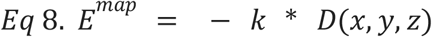

Where, D(x,y,z) is the normalised density value at position x,y,z within the full map.

Following minimisation, the system’s temperature was gradually raised from 0K to 300K in 1K increments. At each increment, the simulation was equilibrated for 1000 steps.

Once the desired temperature was reached, the system was heated from 300 K to 315 K by temperature steps of 1K. Again at each temperature step the system was allowed to advance 1000 steps, before being cooled to 300 K in the same manner. This continued for a defined amount of cycles (4 by default), before yielding the final solutions.

For both minimisation and simulated annealing atoms not within binding sites or forming part of the protein backbones were not minimised/refined. These atoms were kept in place using a harmonic pinning potential whose strength(*k_pin_*) was 500 kJ/mol.

The final solutions were grouped by their RMSD (< 2.0Å), where each group was represented by the solution with the highest ChemEM/ChemDock score.

### Molecular docking experiments and small molecule fitting

For molecular docking experiments smiles strings for the 32 ligands in the benchmark were extracted from the PDB chemical repository. Protein models were preprecessed by removing solvent and ligands. The binding site was defined from the native structure as the centroid of the deposited ligand, a radius of 12.5Å was used for ChemDock experiments and a box size of 12.5Å for docking with AutoDock vina. When using the ChemEM software, protein models were input as pre-processed PDB files, whilst ligands were supplied as a smiles string. For AutoDock vina preprocessed protein files were prepared using the *‘DockPrep’* command in UCSF Chimera. Ligands for AutoDock Vina were prepared by first generating a 3D conformation in RDKit using the SMILE strings. These ligands were then prepared into PDBQT files using the Chimera *‘DockPrep’*.

For multi-ligand docking experiments, the ChemEM software was extended to receive multiple inputs within a single binding site. Ligands were given as individual SMILES strings. Regarding the difference map inputs and centre of binding sites, there are two options, the first was to give a difference map and binding site centroid as an input. The second option (used for experiments reported here) is to give multiple difference map inputs and binding site centroids. The search space for each ligand was then defined from the assigned difference density, with the centre of the ligand binding site for a specific ligand being the centre point of the assigned difference density. To fit multiple ligands into the binding site simultaneously the scoring function was extended such that inter-ligand interactions were accounted for in the same manner as protein-ligand interactions when using the ChemDock score.

To perform multi-ligand docking first ligand density within the difference map was assigned to a given ligand and disconnected difference densities extracted from the whole difference map using a map processing function contained in ChemEM (Figure S2A). Briefly, this command grouped regions of density together where the boundaries were defined as voxels with any adjacent voxel below a given threshold (0.0 by default). The densities assumed to correspond to individual ligands were extracted from the whole map and all voxels in a given disconnected density were vectorised and the centroid for the binding site for that ligand was determined by the centroid of the vectorised points (Figure S2B).

To evaluate the solutions given by the docking/fitting programs the RMSD to the reference structure was calculated using the symmetry-corrected RMSD implemented in the python package SpyRMSD(38). For each experiment the RMSDs given represented the mean minimum RMSD, this was defined as the average of the solutions for each case with the lowest RMSD to the reference structure.

Additionally, the strain energy of ligand conformations was calculated using the method outlined in Gu *et al*. (17). Calculation of the cross-orrelation coefficient was conducted using the python package TEMPy (31), using full maps downloaded from the electron microscopy data bank (1).

## Acknowledgments

We thank Tristan Cragnolini, Ronja Markworth, Mauro Maiorca and the Topf lab for helpful discussions. This work was supported by the Wellcome Trust grant 209250/Z/17/Z and cooperation of Leibniz Institute of Virology (as part of Leibniz ScienceCampus InterACt, funded by the BWFGB Hamburg and the Leibniz Association).

## References

1. C. L. Lawson, et al., EMDataBank unified data resource for 3DEM. Nucleic Acids Res. 44, D396–403 (2016).

2. J. J. Kim, et al., Shared structural mechanisms of general anaesthetics and benzodiazepines. Nature 585, 303–308 (2020).

3. T. Nakane, et al., Single-particle cryo-EM at atomic resolution. Nature 587, 152–156 (2020).

4. M. J. Robertson, G. C. P. van Zundert, K. Borrelli, G. Skiniotis, GemSpot: A Pipeline for Robust Modeling of Ligands into Cryo-EM Maps. Structure 28, 707–716.e3 (2020).

5. J. W. Vant, et al., Flexible Fitting of Small Molecules into Electron Microscopy Maps Using Molecular Dynamics Simulations with Neural Network Potentials. J. Chem. Inf. Model. 60, 2591–2604 (2020).

6. O. Trott, A. J. Olson, AutoDock Vina: improving the speed and accuracy of docking with a new scoring function, efficient optimization, and multithreading. J. Comput. Chem. 31, 455–461 (2010).

7. M. Su, et al., Comparative Assessment of Scoring Functions: The CASF-2016 Update. J. Chem. Inf. Model. 59, 895–913 (2019).

8. D. Vasishtan, M. Topf, Scoring functions for cryoEM density fitting. J. Struct. Biol. 174, 333–343 (2011).

9. J. Locke, et al., Structural basis of human kinesin-8 function and inhibition. Proc. Natl. Acad. Sci. U. S. A. 114, E9539–E9548 (2017).

10. A. Peña, et al., Structure of Microtubule-Trapped Human Kinesin-5 and Its Mechanism of Inhibition Revealed Using Cryoelectron Microscopy. Structure 28, 450–457.e5 (2020).

11. G. Jones, P. Willett, R. C. Glen, A. R. Leach, R. Taylor, Development and validation of a genetic algorithm for flexible docking11Edited by F. E. Cohen. J. Mol. Biol. 267, 727–748 (1997).

12. G. C. P. van Zundert, et al., The HADDOCK2.2 Web Server: User-Friendly Integrative Modeling of Biomolecular Complexes. J. Mol. Biol. 428, 720–725 (2016).

13. A. P. Joseph, et al., Comparing Cryo-EM Reconstructions and Validating Atomic Model Fit Using Difference Maps. J. Chem. Inf. Model. 60, 2552–2560 (2020).

14. J. A. Maier, et al., ff14SB: Improving the Accuracy of Protein Side Chain and Backbone Parameters from ff99SB. J. Chem. Theory Comput. 11, 3696–3713 (2015).

15. Y. Qiu, et al., Development and Benchmarking of Open Force Field v1.0.0—the Parsley Small-Molecule Force Field. J. Chem. Theory Comput. 17, 6262–6280 (2021).

16. O. Korb, T. Stützle, T. E. Exner, An ant colony optimization approach to flexible protein–ligand docking. Swarm Intelligence 1, 115–134 (2007).

17. S. Gu, M. S. Smith, Y. Yang, J. J. Irwin, B. K. Shoichet, Ligand Strain Energy in Large Library Docking. J. Chem. Inf. Model. 61, 4331–4341 (2021).

18. O. Korb, T. Stützle, T. E. Exner, PLANTS: Application of Ant Colony Optimization to Structure-Based Drug Design in Ant Colony Optimization and Swarm Intelligence, (Springer Berlin Heidelberg, 2006), pp. 247–258.

19. M. Su, et al., Structural Basis of the Activation of Heterotrimeric Gs-Protein by Isoproterenol-Bound β1-Adrenergic Receptor. Mol. Cell 80, 59–71.e4 (2020).

20. X. Zhang, et al., Differential GLP-1R Binding and Activation by Peptide and Non-peptide Agonists. Mol. Cell 80, 485–500.e7 (2020).

21. Y. Kim, E. Jeong, J.-H. Jeong, Y. Kim, Y. Cho, Structural Basis for Activation of the Heterodimeric GABAB Receptor. J. Mol. Biol. 432, 5966–5984 (2020).

22. J. Gutiérrez-Fernández, et al., Key role of quinone in the mechanism of respiratory complex I. Nat. Commun. 11, 4135 (2020).

23. J. Zhu, K. R. Vinothkumar, J. Hirst, Structure of mammalian respiratory complex I. Nature 536, 354–358 (2016).

24. S. Pospich, F. Merino, S. Raunser, Structural Effects and Functional Implications of Phalloidin and Jasplakinolide Binding to Actin Filaments. Structure 28, 437–449.e5 (2020).

25. M. Saur, et al., Fragment-based drug discovery using cryo-EM. Drug Discov. Today 25, 485–490 (2020).

26. I. Farabella, et al., TEMPy: a Python library for assessment of three-dimensional electron microscopy density fits. J. Appl. Crystallogr. 48, 1314–1323 (2015).

27. A. P. Joseph, I. Lagerstedt, A. Patwardhan, M. Topf, M. Winn, Improved metrics for comparing structures of macromolecular assemblies determined by 3D electron-microscopy. J. Struct. Biol. 199, 12–26 (2017).

28. R. Wang, X. Fang, Y. Lu, S. Wang, The PDBbind Database: Collection of Binding Affinities for Protein-Ligand Complexes with Known Three-Dimensional Structures. J. Med. Chem. 47, 2977–2980 (2004).

29. M. Wójcikowski, P. Zielenkiewicz, P. Siedlecki, Open Drug Discovery Toolkit (ODDT): a new open-source player in the drug discovery field. J. Cheminform. 7, 26 (2015).

30. E. F. Pettersen, et al., UCSF Chimera--a visualization system for exploratory research and analysis. J. Comput. Chem. 25, 1605–1612 (2004).

31. T. Cragnolini, et al., TEMPy2: a Python library with improved 3D electron microscopy density-fitting and validation workflows. Acta Crystallogr D Struct Biol 77, 41–47 (2021).

32. H. Berman, K. Henrick, H. Nakamura, J. L. Markley, The worldwide Protein Data Bank (wwPDB): ensuring a single, uniform archive of PDB data. Nucleic Acids Res. 35, D301–3 (2007).

33. R. Wang, L. Liu, L. Lai, Y. Tang, SCORE: A New Empirical Method for Estimating the Binding Affinity of a Protein-Ligand Complex. Molecular modeling annual 4, 379–394 (1998).

34. R. Wang, L. Lai, S. Wang, Further development and validation of empirical scoring functions for structure-based binding affinity prediction. J. Comput. Aided Mol. Des. 16, 11–26 (2002).

35. T. Cheng, et al., Computation of octanol-water partition coefficients by guiding an additive model with knowledge. J. Chem. Inf. Model. 47, 2140–2148 (2007).

36. M. Brylinski, Aromatic interactions at the ligand-protein interface: Implications for the development of docking scoring functions. Chem. Biol. Drug Des. 91, 380–390 (2018).

37. P. Eastman, V. S. Pande, OpenMM: A Hardware Independent Framework for Molecular Simulations. Comput. Sci. Eng. 12, 34–39 (2015).

38. R. Meli, P. C. Biggin, spyrmsd: symmetry-corrected RMSD calculations in Python. J. Cheminform. 12, 49 (2020).

