## Supplemental File for "ChemEM: flexible docking of small molecules in Cryo-EM structures using difference maps"

### Supplementary Figures

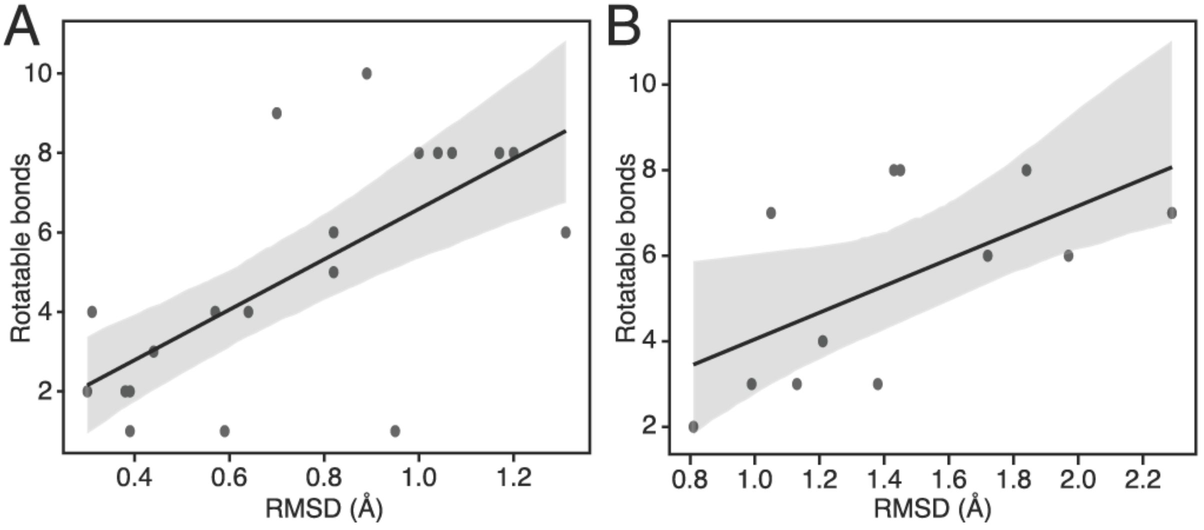


**Figure S1.** The correlation between the number of rotatable bonds within a small molecule and the RMSD of the best solutions to the PDB deposited structure for the high- (**A**) and low- (**B**) resolution benchmarks.

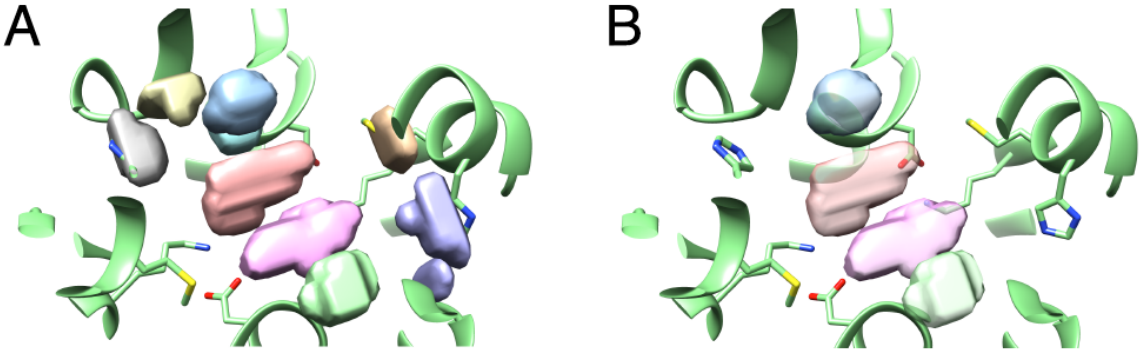


**Figure S2**. The difference mapping procedure for multiple ligand fitting. **A**. The local difference map within the binding site after disconnected densities (various colours) were identified by ChemEM. **B**. The final difference map once disconnected densities that correspond to protein atoms were removed. Each disconnected density (in blue, peach, pink and green) corresponded to an individual ligand within the binding site, and was used for binding site centroid calculations.

### Supplementary Tables

**Table S1**. 5 fold validation results from training the ChemDock scoring function.

| Group | Pearson Correlation (train) | Pearson Correlation (test) |
|---|---|---|
| 1 | 0.57 | 0.58 |
| 2 | 0.57 | 0.57 |
| 3 | 0.56 | 0.60 |
| 4 | 0.57 | 0.50 |
| 5 | 0.56 | 0.61 |

**Table S2**. Scoring function weights for each term in the ChemDock scoring function.

| Term | Weight | Term | Weight |
|---|---|---|---|
| HBD | -0.079 | ArmD | -0.508 |
| HBA | -0.041 | ArmA | 0.534 |
| VG1 | 0.0087 | $VG1_{intra}$ | -0.0062 |
| VG2 | 0.0016 | $VG2_{intra}$ | -0.0012 |
| VS | -0.0819 | $VS_{intra}$ | -0.016 |
| LogP | 0.0077 | Bias | 3.476 |

**Table S3.** The PDB ID, Ligand ID, EMBD ID, and resolutions of the high and low-resolution benchmarks.

| PDB | Ligand | EMBD | Resolution | PDB | ligand | EMDB | Resolution |
|---|---|---|---|---|---|---|---|
| High Resolution | | | | | | | |
| 6X40 | RI5 | 22038 | 2.86 | 6X3T | PFL | 22032 | 2.55 |
| 6TTE | PTQ | 10574 | 2.2 | 6KPF | 8D0 | 0744 | 2.9 |
| 6UDP | IMP | 20742 | 2.95 | 6TTI | NXE | 10577 | 2.5 |
| 6TTQ | FBP | 10584 | 2.7 | 6X1A | UK4 | 21994 | 2.5 |
| 6A95 | 9SR | 6997 | 2.6 | 7JJO | 5FW | 22357 | 2.6 |
| 6OAX | AGS | 20004 | 2.9 | 6VFX | ATP | 21194 | 2.9 |
| 6PEQ | ZK1 | 20330 | 2.97 | 6NYY | ANP | 0552 | 3.0 |
| 6UZ8 | R0D | 20953 | 2.84 | 7C7Q | 2C0 | 30300 | 3.0 |
| 6UQE | ADP | 20845 | 3.0 | 7CFM | FWX | 30344 | 3.0 |
| 6OO3 | 6EU | 20143 | 2.9 | 6X3X | DZP | 22036 | 2.92 |
| Low Resolution | | | | | | | |
| 5OAF | ADP | 3773 | 4.06 | 5W3J | GTP | 8757 | 4.0 |
| 6R4O | FOK | 4721 | 4.2 | 6T24 | 9ZK | 10366 | 3.7 |
| 6ZIY | FMN | 11231 | 4.25 | 6WHV | QGP | 21677 | 4.05 |
| 7CUM | 2BV | 30472 | 3.52 | 6IP2 | ATP | 9698 | 3.7 |
| 6K42 | CZX | 9912 | 4.1 | 5WEL | ZK1 | 8820 | 4.4 |
| 6U8S | IMP | 20691 | 3.14 | 6N57 | 1N7 | 0348 | 3.7 |

**Table S4.** The RMSD, CCC, TEU of the best final solutions for AutoDock Vina, ChemDock and ChemEM, for the high-resolution benchmark.

| | Deposited | | AutoDock Vina | | | ChemDock | | | ChemEM | | |
|---|---|---|---|---|---|---|---|---|---|---|---|
| PDB | CCC | TEU | RMSD | CCC | TEU | RMSD | CCC | TEU | RMSD | CCC | TEU |
| 6oo3 | 0.017 | 16.69 | 1.86 | 0.013 | 16.27 | 1.39 | 0.014 | 17.46 | 0.7 | 0.016 | 13.644 |
| 6tti | 0.011 | 1.12 | 3.7 | 0.01 | 1.75 | 3.41 | 0.009 | 0.77 | 0.39 | 0.011 | 1.15 |
| 7cfm | 0.034 | 10.74 | 3.16 | 0.014 | 7.2 | 1.39 | 0.029 | 9.81 | 1.17 | 0.034 | 6.78 |
| 6nyy | 0.035 | 7.34 | 4.57 | 0.028 | 7.36 | 2.33 | 0.030 | 3.10 | 1.2 | 0.033 | 5.777 |
| 6vfx | 0.024 | 7.73 | 4.43 | 0.016 | 8.77 | 1.05 | 0.022 | 5.53 | 1.07 | 0.024 | 7.222 |
| 6kpf | 0.019 | 18.9 | 1.39 | 0.018 | 25.69 | 1.19 | 0.018 | 21.97 | 1.00 | 0.021 | 13.39 |
| 6a95 | 0.024 | 1.96 | 0.56 | 0.022 | 1.06 | 1.01 | 0.021 | 0.971 | 0.59 | 0.023 | 0.567 |
| 7c7q | 0.026 | 6.14 | 0.54 | 0.026 | 6.23 | 0.70 | 0.025 | 2.87 | 0.31 | 0.026 | 4.684 |
| 6udp | 0.021 | 4.87 | 1.38 | 0.015 | 4.46 | 1.41 | 0.013 | 4.73 | 0.57 | 0.021 | 1.879 |
| 6x1a | 0.032 | 13.36 | 1.26 | 0.026 | 13.22 | 2.09 | 0.017 | 13.41 | 0.89 | 0.03 | 11.178 |
| 6uz8 | 0.022 | 1.81 | 4.76 | 0.015 | 2.58 | 1.25 | 0.018 | 1.94 | 0.3 | 0.021 | 2.102 |
| 6x3t | 0.015 | 0.62 | 0.85 | 0.013 | 0.34 | 0.78 | 0.014 | 1.04 | 0.38 | 0.015 | 0.32 |
| 7jjo | 0.036 | 3.91 | 0.46 | 0.036 | 2.27 | 0.72 | 0.036 | 2.18 | 0.64 | 0.037 | 1.202 |
| 6x40 | 0.01 | 0.65 | 1.13 | 0.01 | 1.18 | 1.76 | 0.009 | 1.58 | 0.95 | 0.009 | 0.283 |
| 6uqe | 0.018 | 5.35 | 1.41 | 0.018 | 4.1 | 1.29 | 0.017 | 4.58 | 1.31 | 0.018 | 4.155 |
| 6peq | 0.029 | 3.86 | 0.85 | 0.028 | 3.82 | 1.36 | 0.027 | 2.42 | 0.44 | 0.03 | 3.052 |
| 6ttq | 0.03 | 10.37 | 1.53 | 0.019 | 8.51 | 1.57 | 0.018 | 9.23 | 0.82 | 0.031 | 5.499 |
| 6x3x | 0.013 | 0.97 | 1.06 | 0.013 | 0.916 | 1.68 | 0.01 | 2.08 | 0.39 | 0.015 | 1.381 |
| 6tte | 0.013 | 5.71 | 1.03 | 0.011 | 7.19 | 1.06 | 0.011 | 3.60 | 0.82 | 0.013 | 9.223 |
| 6oax | 0.02 | 7.43 | 2.62 | 0.015 | 7.9 | 1.45 | 0.018 | 8.89 | 1.04 | 0.019 | 3.803 |
| mean | 0.023 | 6.47 | 1.93 | 0.018 | 6.53 | 1.44 | 0.019 | 5.90 | 0.75 | 0.22 | 4.86 |

**Table S5.** The RMSD, CCC, TEU of the best final solutions for AutoDock Vina, ChemDock and ChemEM, for the low-resolution benchmark.

| | Deposited | | HR* | AutoDock Vina | | | ChemDock | | | ChemEM | | |
|---|---|---|---|---|---|---|---|---|---|---|---|---|
| PDB | CCC | TEU | TEU | RMSD | CCC | TEU | RMSD | CCC | TEU | RMSD | CCC | TEU |
| 5oaf | 0.039 | 1.55 | 5.35 | 1.07 | 0.038 | 5.81 | 1.92 | 0.035 | 7.34 | 1.72 | 0.037 | 1.85 |
| 6r4o | 0.025 | 6.97 | 5.53 | 1.73 | 0.024 | 7.72 | 1.46 | 0.024 | 5.71 | 1.13 | 0.025 | 2.02 |
| 6ziy | 0.005 | 11.37 | 12.24 | 1.12 | 0.005 | 11.48 | 1.22 | 0.005 | 8.57 | 1.05 | 0.005 | 11.21 |
| 7cum | 0.029 | 6.76 | 6.76 | 2.059 | 0.029 | 8.17 | 3.33 | 0.02 | 11.56 | 1.84 | 0.031 | 10.61 |
| 6k42 | 0.021 | 0.51 | 0.49 | 0.44 | 0.023 | 0.27 | 0.68 | 0.022 | 0.54 | 0.81 | 0.021 | 0.84 |
| 6u8s | 0.03 | 6.14 | 6.14 | 2.92 | 0.014 | 6.43 | 1.97 | 0.026 | 5.77 | 1.21 | 0.027 | 5.24 |
| 5w3j | 0.011 | 6.04 | 7.02 | 1.80 | 0.01 | 7.72 | 0.98 | 0.01 | 9.19 | 1.45 | 0.011 | 7.93 |
| 6t24 | 0.027 | 6.67 | 9.19 | 7.10 | 0.014 | 18.50 | 3.01 | 0.02 | 12.71 | 2.29 | 0.027 | 6.97 |
| 6whv | 0.017 | 5.71 | 6.45 | 1.74 | 0.013 | 5.65 | 1.62 | 0.014 | 16.32 | 1.97 | 0.016 | 5.11 |
| 6ip2 | 0.017 | 5.69 | 5.69 | 2.21 | 0.014 | 5.83 | 1.93 | 0.014 | 7.37 | 1.43 | 0.017 | 7.34 |
| 5wel | 0.038 | 4.61 | 2.21 | 1.22 | 0.035 | 4.05 | 1.22 | 0.035 | 6.5 | 0.99 | 0.038 | 2.91 |
| 6n57 | 0.012 | 5.65 | 4.86 | 7.25 | 0.011 | 10.54 | 1.24 | 0.01 | 8.3 | 1.38 | 0.013 | 5.36 |
| mean | 0.023 | 5.64 | 5.99 | 2.55 | 0.019 | 7.68 | 1.72 | 0.02 | 8.32 | 1.44 | 0.022 | 5.61 |

**Table S6.** Rank data for ChemDock solutions with the best RMSD in the high-resolution benchmark.

| PDB | Rank | *n* solutions | Rank (%) | Correct identified |
|---|---|---|---|---|
| **6oo3** | 1 | 183 | 0.55 | True |
| **6tti** | 13 | 48 | 27.08 | False |
| **7cfm** | 89 | 127 | 70.08 | True |
| **6nyy** | 12 | 170 | 7.06 | False |
| **6vfx** | 5 | 169 | 2.96 | True |
| **6kpf** | 36 | 144 | 25.0 | True |
| **6a95** | 1 | 43 | 2.33 | True |
| **7c7q** | 6 | 66 | 9.09 | True |
| **6udp** | 13 | 86 | 15.12 | True |
| **6x1a** | 37 | 209 | 17.7 | False |
| **6uz8** | 6 | 88 | 8.82 | True |
| **6x3t** | 1 | 26 | 3.85 | True |
| **7jjo** | 17 | 77 | 22.09 | True |
| **6x40** | 7 | 22 | 21.21 | True |
| **6uqe** | 7 | 152 | 4.61 | True |
| **6peq** | 9 | 82 | 10.98 | True |
| **6ttq** | 3 | 94 | 3.19 | True |
| **6x3x** | 13 | 45 | 28.89 | True |
| **6tte** | 25 | 103 | 24.27 | True |
| **6oax** | 1 | 164 | 0.61 | True |

**Table S7.** Rank data for ChemDock solutions with the best RMSD in the low-resolution benchmark.

| PDB | Rank | *n* solutions | Rank (%) | Correct identified |
|---|---|---|---|---|
| **5oaf** | 7 | 147 | 4.76 | True |
| **6r4o** | 4 | 55 | 7.27 | True |
| **6ziy** | 3 | 152 | 1.97 | True |
| **7cum** | 73 | 203 | 35.96 | False |
| **6k42** | 7 | 68 | 10.29 | True |
| **6u8s** | 27 | 88 | 30.68 | True |
| **5w3j** | 5 | 189 | 2.65 | True |
| **6t24** | 15 | 158 | 9.49 | False |
| **6whv** | 16 | 154 | 10.39 | True |
| **6ip2** | 62 | 180 | 34.44 | True |
| **5wel** | 1 | 77 | 1.3 | True |
| **6n57** | 56 | 232 | 24.14 | True |

**Table S8.** A Table of average Pearson correlation coefficients between the MI/CCC of simulated difference maps and RMSD of the CASF-2016 decoy sets at resolutions between 2.5 and 8.5Å.

| Resolution (Å) | 2.5 | 3.0 | 3.5 | 4.0 | 4.5 | 5.0 | 5.5 | 6.0 | 6.5 | 7.0 | 7.5 | 8.0 | 8.5 |
|---|---|---|---|---|---|---|---|---|---|---|---|---|---|
| MI | -0.802 | -0.805 | -0.808 | -0.812 | -0.812 | -0.812 | -0.813 | -0.814 | -0.813 | -0.812 | -0.810 | -0.810 | -0.809 |
| CCC | -0.792 | -0.789 | -0.787 | -0.787 | -0.782 | -0.777 | -0.773 | -0.772 | -0.768 | -0.766 | -0.762 | -0.761 | -0.759 |
| *p** | 0.41 | 0.24 | 0.14 | 0.067 | 0.038 | 0.017 | 0.008 | 0.005 | 0.0035 | 0.003 | 0.002 | 0.002 | 0.0016 |

** P values are calculated with a Student's T-test between the results of individual groups of MI and CCC at each resolution.*

**Table S9.** Rank data for ChemEM solutions with the best RMSD in the high-resolution benchmark.

| **PDB** | **Rank** | ***n* solutions** | **Rank (%)** | **Correct identified** |
|---|---|---|---|---|
| **6oo3** | 11 | 211 | 5.21 | True |
| **6tti** | 13 | 61 | 21.31 | True |
| **7cfm** | 12 | 176 | 6.82 | True |
| **6nyy** | 2 | 157 | 1.27 | True |
| **6vfx** | 36 | 193 | 18.65 | True |
| **6kpf** | 1 | 115 | 0.87 | True |
| **6a95** | 1 | 43 | 2.33 | True |
| **7c7q** | 1 | 63 | 1.59 | True |
| **6udp** | 2 | 90 | 2.22 | True |
| **6x1a** | 15 | 250 | 6.0 | True |
| **6uz8** | 4 | 86 | 4.65 | True |
| **6x3t** | 1 | 36 | 2.78 | True |
| **7jjo** | 1 | 67 | 1.49 | True |
| **6x40** | 1 | 34 | 2.94 | True |
| **6uqe** | 6 | 137 | 4.38 | True |
| **6peq** | 2 | 76 | 2.63 | True |
| **6ttq** | 2 | 90 | 2.22 | True |
| **6x3x** | 1 | 36 | 2.78 | True |
| **6tte** | 1 | 104 | 0.96 | True |
| **6oax** | 1 | 183 | 0.55 | True |

**Table S10.** Rank data for ChemEM solutions with the best RMSD in the low-resolution benchmark.

| PDB | Rank | *n* solutions | Rank (%) | Correct identified |
|---|---|---|---|---|
| **5oaf** | 1 | 129 | 0.78 | True |
| **6r4o** | 2 | 47 | 4.26 | True |
| **6ziy** | 26 | 146 | 17.81 | True |
| **7cum** | 58 | 224 | 25.89 | True |
| **6k42** | 11 | 65 | 16.92 | True |
| **6u8s** | 10 | 86 | 11.63 | True |
| **5w3j** | 36 | 172 | 20.93 | True |
| **6t24** | 7 | 130 | 5.38 | False |
| **6whv** | 1 | 118 | 0.85 | True |
| **6ip2** | 11 | 169 | 6.51 | True |
| **5wel** | 1 | 62 | 1.61 | True |
| **6n57** | 28 | 242 | 11.57 | True |

**Table S11.** The percent of ChemEM cases where a correct solution was identified for the high- and low-resolution benchmarks using a global k of 25, 50, and 75.

| K | RMSD ≤ 1.0Å (%) | RMSD ≤ 1.5Å (%) | RMSD ≤ 2.0Å (%) |
|---|---|---|---|
| **High resolution** | | | |
| **25** | 50 | 80 | 95 |
| **50** | 60 | 85 | 95 |
| **75** | 75 | 100 | 100 |
| **Low resolution** | | | |
| **25** | 16.6 | 66.6 | 91.6 |
| **50** | 8.3 | 25.0 | 83.3 |
| **75** | 16.6 | 33.3 | 91.6 |
